# Functional single cell proteomic profiling of cells with abnormal DNA damage response dynamics

**DOI:** 10.1101/2021.10.13.464241

**Authors:** Pin-Rui Su, Li You, Cecile Beerens, Karel Bezstarosti, Jeroen Demmers, Martin Pabst, Roland Kanaar, Cheng-Chih Hsu, Miao-Ping Chien

**Author notes:** co-senior author.

## Abstract

Tumor heterogeneity is an important source of cancer therapy resistance. Single cell proteomics has the potential to decipher protein content leading to heterogeneous cellular phenotypes. Single-Cell ProtEomics by Mass Spectrometry (SCoPE-MS) is a recently developed, promising, unbiased proteomic profiling techniques, which allows profiling several tens of single cells for >1000 proteins per cell. However, a method to link single cell proteomes with cellular behaviors is needed to advance this type of profiling technique. Here, we developed a microscopy-based functional single cell proteomic profiling technology, called FUNpro, to link the proteome of individual cells with phenotypes of interest, even if the phenotypes are dynamic or the cells of interest are sparse. FUNpro enables one i) to screen thousands of cells with subcellular resolution and monitor (intra)cellular dynamics using a custom-built microscope, ii) to real-time analyze (intra)cellular dynamics of individual cells using an integrated cell tracking algorithm, iii) to promptly isolate the cells displaying phenotypes of interest, and iv) to single cell proteomically profile the isolated cells. We applied FUNpro to proteomically profile a newly identified small subpopulation of U2OS osteosarcoma cells displaying an abnormal, prolonged DNA damage response (DDR) after ionizing radiation (IR). With this, we identified PDS5A and PGAM5 proteins contributing to the abnormal DDR dynamics and helping the cells survive after IR.

---

A high degree of cellular heterogeneity underlies biological processes like tumorigenesis or differentiation. In the past several years, single cell profiling technologies have revolutionized biological and biomedical research into rare cells or subpopulations of cells^1,2^. Single cell RNA or DNA sequencing technology has developed much more rapidly than single cell proteomics^3^. However, to gain mechanistic understanding of cellular processes like the DNA damage response (DDR), measuring and quantifying the effector/active molecules, namely the proteins, is imperative^3^. SCoPE-MS is a recently developed, antibody-independent single cell proteomics technique^4^; this approach has gradually become popular because it is an unbiased proteomic profiling method (antibody-independent) and enables identification of >1000 proteins in single cells. Cell sorting is often applied before SCoPE-MS, and cell selection based on static features has advanced rapidly in recent years^5^. However, a method to link measured proteomes of single cells to more interesting cellular, intracellular and intercellular dynamics (for example migration, longitudinal protein dynamics, multicell interaction), static features and combinations of both, does not yet exist and would significantly expand on the applications for single cell proteomics, allowing the investigation of novel mechanistic questions.

Here, we introduce a technology, called <u>fun</u>ctional single cell <u>pro</u>teomic profiling (FUNpro, Figure 1a), that 1) enables screening a population containing a large quantity of cancer cells (> 10^3^) with high spatiotemporal resolution via a custom-built Ultrawide Field-of-view Optical (UFO) microscope, 2) allows real-time identifying cells with different (intra)cellular dynamics via an integrated automatic cell tracking algorithm, and 3) permits separating different phenotypes of cells with selective photolabeling of desired cells followed by cell sorting and single cell proteomic profiling. With FUNpro, we can stratify or pre-select cells ahead of time based on any microscopically observable cellular and/or intracellular behaviors before subjecting to single cell proteomic measurements.

**Figure 1.**
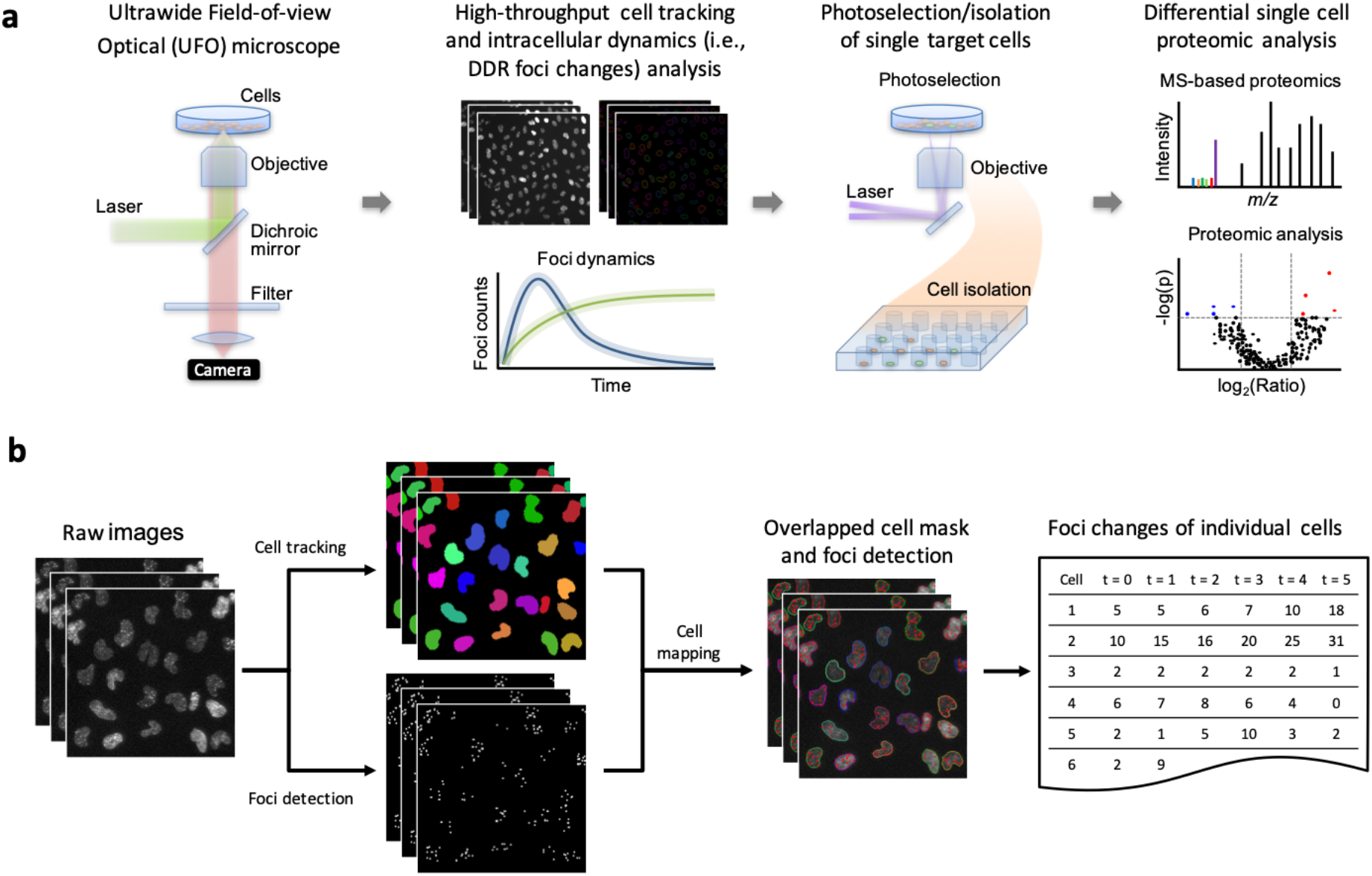
Outline of the developed functional single cell proteomic profiling (FUNpro) pipeline. (a) FUNpro pipeline: cells were high-throughput screened under the UFO microscope followed by real-time cell tracking and intracellular dynamics analysis to identify cells of interest; desired cells were then selectively photolabelled followed by cell sorting before being subjected to single cell proteomic measurements and analysis. (b) Schematic of high-throughput identification and selection of target cells via an automated image processing and analysis algorithm.

An example of a heterogeneous phenomenon that can be investigated using this technique is DNA damage repair (DDR). DDR, triggered by DNA breaks or DNA double-strand breaks (DSBs), leads to a series of DDR proteins to assemble and act on damaged DNA sites to maintain genomic integrity^6–8^. One of the key DDR proteins, tumor suppressor p53-binding protein 1 (53BP1), accumulates and forms oligomers (53BP1 foci) on DSBs to regulate DNA repair and has been used as a DDR indicator in response to DSBs^6–8^. Unrepaired or misrepaired DSBs often lead to mutations or chromosomal rearrangements that can result in cell death or oncogenic transformation that can promote tumor progression and evolution^9^. In addition, efficient DNA damage repair, caused by among others overactive or overexpressed DDR proteins, can lead to cancer cell survival in response to treatment^9^. These various scenarios regarding DNA damage response/repair mechanisms can contribute to tumor heterogeneity and lead to different cell fates (cell death or tumor progression); hence, it is crucial to decipher causative underlying mechanisms of heterogeneous dynamics of DDR.

As a proof-of-concept, we applied FUNpro to profile and investigate a subpopulation of U2OS cells displaying abnormal DDR induced by ionizing radiation (IR). We used U2OS cells expressing 53BP1-mScarlet as a model system and monitored IR-induced DDR dynamics via 53BP1 foci changes, imaged through clustered mScarlet-fluorescence. To screen a large quantity of cells and identify subpopulations of cells with different intracellular 53BP1 foci changes over time, we implemented the UFO microscope^10^ in the FUNpro pipeline (Figure 1a) to image thousands of cells with 0.8 μm spatial resolution, sufficient to resolve nuclear DDR foci. We developed and implemented an automatic DDR foci tracking algorithm (Methods, Figure 1b, S1-3) in the modified Tracking with Gaussian Mixture Model (mTGMM^10^) cell tracking algorithm, to quantify changes in DDR foci while tracking cellular movement in a real-time fashion. We tracked individual cell migration, division and intracellular dynamics for thousands of cells (~57,000 cells/min) to register foci dynamics for individual cells over hundreds of image frames (24 hours of imaging; Figure 1b, S1-3). We applied a phototagging technique (Methods) to selectively photolabel cells of interest for cell sorting with Fluorescence Activated Cell Sorting (FACS). Finally, SCoPE-MS was applied to quantify the proteins of single cells and a differential protein analysis was computed by comparing the proteins from the cells of interest with the control cells.

After irradiating U2OS-53BP1-mScarlet cells with 2 Gy IR, cells were immediately subjected to imaging by UFO for data acquisition and imaging for 24 hrs (3 min/frame). We then applied the integrated cell and DDR foci tracking algorithm (Methods, Figure 1b, S1-3) to analyze the 24 hrtime lapse movie of U2OS-53BP1-mScarlet cells (Figure 2a). After analyzing ~10^3^ irradiated cells, we identified two different groups of cells based on the dynamics of DDR foci formation (53BP1-mScarlet foci) (Figure 2b,c). One group (Group 1) had a peak amount of DDR foci at 4-6 hr after radiation, immediately followed by decay in foci number at 6 hr (we call this up-&-down foci trend or normal DDR dynamics, based on published observations^11–13^). The other group (Group 2) had rising DDR foci counts at 4-6 hr without decaying until at least 24 hr (we call this rising foci trend or abnormal DDR dynamics) (Figure 2b). The IR-induced DDR dynamics shown in Group 1 is the main phenotype reported previously^11–13^, whereas Group 2 has not been reported before, a phenotype which was only revealed after the cluster analysis from analyzing a large quantity of cells (Figure 2). For the cells showing DDR foci changes, 90.4% of the cells belonged to Group 1 (up-&-down foci trend, Figure 2b) and 9.6% to Group 2 (rising foci trend, Figure 2b). In addition, these two distinct DDR foci dynamic phenotypes were not observed in the absence of IR (Figure S4), indicating that the phenotypes were associated with IR-induced DDR. Given that DDR foci detection has so far mainly been performed using confocal microscopy, we confirmed that the foci detected by UFO show a high correlation (Spearman’s correlation ρ = 0.94, Figure S5c) with the foci detected by confocal microscopy (with size larger than 1 μm in diameter and signal-to-noise (SNR) higher than 2.0) (Figure S5). To further investigate the Group 2 cells, which displayed abnormal, prolonged IR-induced DDR response, we phototagged and separated Group 2 cells and control Group 1 cells (Figure 3a) and performed SCoPE-MS^4^ (Figure 3b, Figure S6). We conducted three independent experiments and collected 40 Group 1 (up-&-down foci trend) and 40 Group 2 (rising foci trend) cells in total after FACS sorting. After annotating cells for Group 1 and Group 2, we performed a differential expression analysis (Figure 3c) and identified a set of differentially expressed proteins (p ≤ 0.05) in Group 2 cells (rising DDR foci). In addition, we implemented a cell cycle scoring analysis from Tirosh *et al.*^14^ and found that both groups of cells had a similar amount of cells at G1, S and G2/M phases (Methods, Figure S7), indicating that these two phenotypes were not caused by cell cycle effects. Amongst the differentially expressed proteins of Group 2 cells, two proteins in particular, sister chromatid cohesion protein PDS5 homolog A (PDS5A) and mitochondrial phosphoglycerate mutase/protein phosphatase (PGAM5), were identified (Figure 3b, c). We validated the discovery of these two proteins, PDS5A and PGAM5, with immunofluorescence staining (Figure 3d). PDS5A has been reported to be a crucial player to protect DNA replication forks^15^, thereby maintaining genome stability from DNA breaks, and the PGAM5 enzyme has been proven to linearly correlate with chemotherapy resistance and preventing apoptosis^16^. Based on this discovery, we suspected that Group 2 cells displaying the rising DDR foci trend after radiation bore more severe DNA damage (more γH2AX foci; Figure S8) than Group 1 cells and were in the process of rescuing themselves from apoptosis and still survived after 2 days of IR (Figure S9).

**Figure 2.**
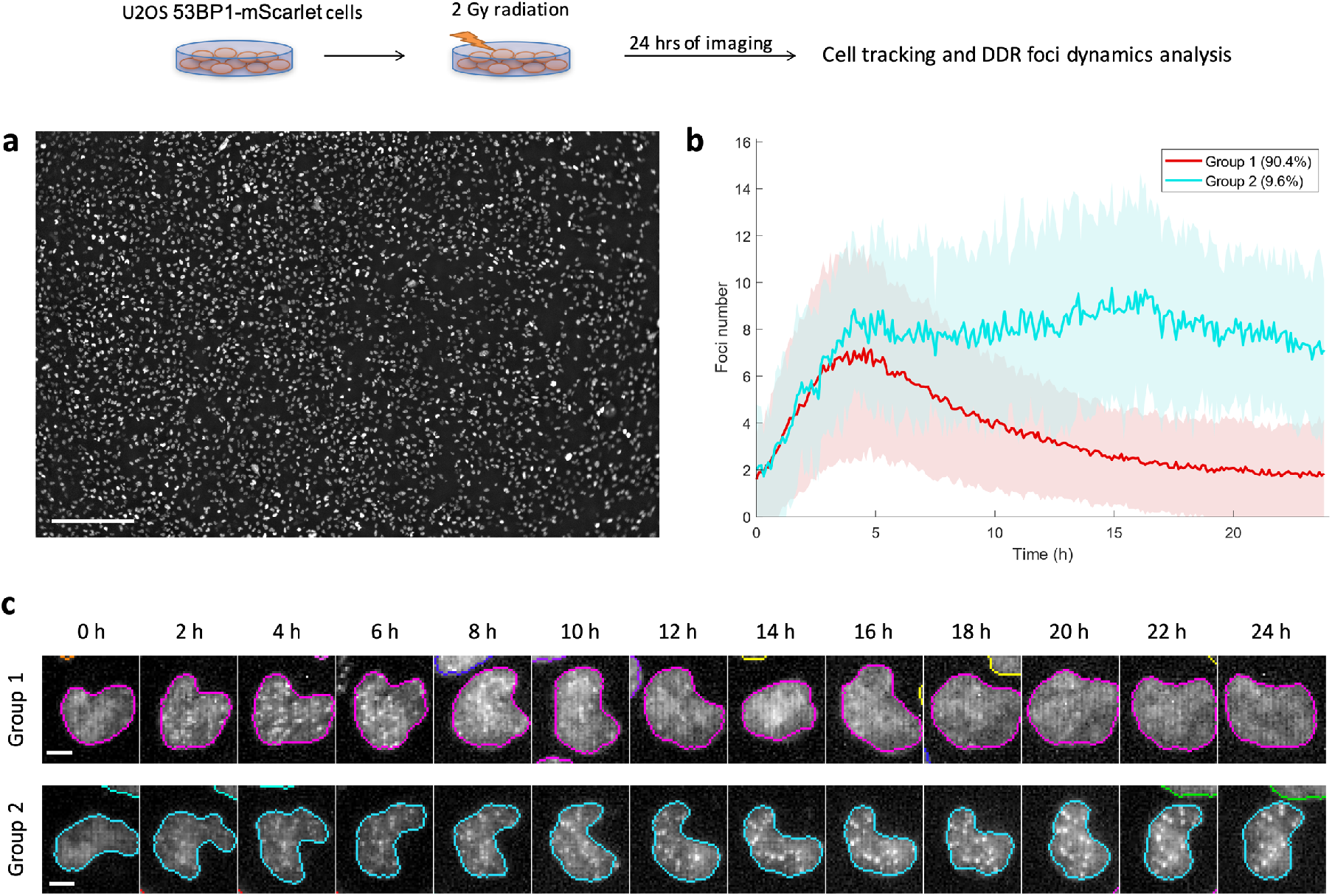
53BP1 foci dynamics tracking (U2OS-53BP1-mScarlet cells) for 24 hrs after 2 Gy irradiation under the UFO microscope. (a) A representative UFO microscopic image of U2OS-53BP1-mScarlet cells post-irradiation. Scale bar = 500 μm. N = ~3000 cells. (b) 53BP1 foci dynamics of two groups of cells over the course of a day. Solid lines represent the average foci number trend from all the cells of each group. The background, light color areas represent the standard deviations of the trends. (c) The zoomed-in images showed the foci changes of representative Group 1 and Group 2 cells after 1 day of IR. Scale bar = 10 μm.

In conclusion, we demonstrated a method to screen a large quantity of individual cells, monitor their cellular and intracellular dynamics, and real-time identify cells of interest displaying desired (intra)cellular dynamics followed by single cell quantitative proteomics. With this, we identified a new subpopulation of U2OS cells displaying abnormal DDR dynamics after IR (rising DDR foci trend) compared to the majority of the cell population (up-&-down foci trend). We also identified PDS5A and PGAM5 proteins as contributing to this cellular phenotype and cell survival after IR. With FUNpro, we can prwe introduce a technology, cae-select individual cells based on any microscopically observable features or characteristics, enrich the quantity of the desired cells and link phenotypes of interest to their proteome. The technology has the potential to investigate causative molecular mechanisms of cells displaying different phenotypes, even if the cells are sparse or dynamic.

## Methods

### Cell culture

Human bone osteosarcoma cell line U2OS was stably transfected with PB-mScarlet-53BP1 (a gift from Hanny Odijk) using Lipofectamine 3000 (Thermo Fisher), under puromycin selection (1 μg/ml). The stable cells were cultured in DMEM supplemented with 10% fetal bovine serum and 1% penicillin/streptomycin in a 37°C incubator under 5% CO_2_.

### Ultrawide field-of-view optical microscope (UFO)

The ultrawide field-of-view optical microscope (UFO) is a custom-built microscope and has been described in detail previously^10^. Here, we upgraded the system to have a higher spatial resolution by implementing an objective with a large field-of-view (FOV) and relatively high numerical aperture (Olympus MVP Plan Apochromat 1x, 0.5 NA) in conjunction with a large chip-size cMOS camera with small pixel-size (Grasshopper3, FLIR, 4096 × 3000 pixels, 3.25 μm/pixel). UFO provides a 3.65 × 2.83 mm FOV with 0.8 μm/pixel spatial resolution and 30 ms/frame temporal resolution.

Illumination source was provided by CW laser lines including 405 nm (MDL-HD-405/2W, CNI), 532 nm (MGL-FN-532/1500mW, CNI) and 637 nm (MDL-MD-637/1.3W, CNI). The lasers were modulated, in wavelength-selection, temporality and intensity, by an acousto-optic tunable filter (Gooch & Housego). Fluorescence was filtered through a custom-designed 2” tri-band emission filter (Od6avg: 400-465 / 527-537 / 632-642 / 785-1300 nm, Alluxa), where filter switching is not needed.

### Cell preparation, radiation and live imaging on UFO microscope

Prior to imaging, U2OS 53BP1-mScarlet cells were seeded in FluoroBrite DMEM supplemented with 10% fetal bovine serum and 1% penicillin/streptomycin at 500k in a 35 mm plastic dish with a 10 mm glass bottom (Cellvis) 1 day in advance. The glass-bottom dish was pre-coated with 0.2% gelatin. The cells were cultured in a 37°C incubator under 5% CO_2_. The mini-incubator at the UFO microscope was equilibrated at 37°C and 5% CO2 prior to imaging. For subsequent phototagging, the cells were pre-incubated with 30 μM photoactivable Janelia Fluor 646 (Bio-Techne) for 15 min, which can be photoactivated by 405 nm laser and visualized by 637 nm laser (λ_ex_: 650 nm; λ_em_: 664 nm). The stained cells were rinsed and incubated in the original medium. Time-lapse movie was recorded by 532 nm laser with interval time of 3 min for 1 hr before ionizing irradiation. After that, the cells were irradiated with a radiation dose of 2 Gy at a dose-rate of 1.67 Gy/min (RS320, Xstrahl Medical & Life Sciences). The cells were put back in the mini-incubator at UFO immediately after radiation and registered back to the previous position. The cells were then recorded by 532 nm laser illumination with interval time of 3 min for 24 hr.

### Automatic foci detection analysis

The foci detection algorithm is a modified local comparison method^17^ (Figure S2). It first generated a reference image from maximum intensity projection of four convoluted images (the raw image was filtered with four different Gaussian-like filter kernels; Figure S2a,b), and then created a normalized reference image by dividing the reference image with a sensitivity factor α (Figure S2c). The sensitivity factor α (0.31) and the radius of filter size (3 pixels) were determined based on the highest F score by sweeping the value of sensitivity factor α from 0 to 1 and the filter size from 0 to 20 pixels, respectively. The foci logical (binary) image was then calculated by Boolean expression (true: when the normalized reference image is greater than the original image; Figure S2a). The detected foci located within a nuclear boundary were registered to the designated nuclei. The number of foci were calculated and plotted over time. Euclidean distance-based hierarchical clustering was used to classify clusters with different trends of foci dynamics. The integrated foci and cell tracking analysis can output the coordinates (in pixels) of target cells with desired features, for instance cells with rising DDR foci. Cells displaying up & down DDR foci dynamics trend were classified as Group 1; cells with rising DDR foci trend were classified as Group 2.

### Automatic cell tracking analysis

To reduce interference from foci when tracking individual cell nuclei, we applied the foci detection algorithm to identify the locations (pixels) of individual foci within each cell nucleus from raw images and removed and replaced them with their neighboring pixels before smoothing the inhomogeneous background within nuclei. We then pre-processed the image using top-hat filtering, local comparison and selection, local brightness adjustment and edge-preserving smoothing to improve accuracy and sensitivity of cell segmentation and tracking. After the foci-dedicated pre-processing, we then applied the mTGMM algorithm for cell segmentation and tracking: we first applied watershed thresholding to find nuclei foreground pixels. The connected foreground pixels were grouped as a single superpixel. All superpixels were trimmed by local Otsu’s thresholding. If more than two superpixels were still connected after thresholding, then they would be grouped together as one. Each final superpixels were fit into Gaussian mixture models (GMMs) separately. With this approach, individual cell nuclei were modeled as GMMs and can be accurately segmented with high precision and recall rate (Precision: 98.1%, Recall: 97.7%, F score: 97.9%, IoU: 95.8%).

The cell tracking was done by forwarding every GMM from time point *t* to *t* + 1 using Bayesian inference, which is, comparing central position, shape, and overall intensity. After cells were tracked, information of cellular characteristics and dynamics including nuclear properties (central position, area, orientation, circularity and intensity), cell division and cell lineage were exported to a feature table. Dividing or apoptotic cells were filtered out during the analysis to perform the Euclidean distance-based hierarchical clustering.

We then applied the integrated foci detection and mTGMM tracking algorithms to simultaneously track cellular movement (with tracking accuracy: 97.7%) while monitoring DDR foci dynamics within each cell nucleus and maintain the processing speed of ~57,000 cells/min. Dividing or dead cells were filtered out during the analysis before performing the Euclidean distance-based hierarchical clustering. All images were processed by MATLAB and C++.

### Target cell photolabeling and isolation

The coordinates (in pixels) of cells of interest identified from the image analysis were then uploaded to the program, which then controlled a pair of galvo mirrors (Cambridge Technology) and steered the selective illumination pattern of 405 nm laser (3 J/cm^2^) onto the cells of interest. Upon 2 s selective illumination, the phototagging reagent inside of the cells under illumination was photoactivated. The photoactivated cells can be invariably visualized by 637 nm laser (λ_ex_: 650 nm; λ_em_: 664 nm), while other cells remain dark. The photoactivated cells can then be isolated (after trypsinization) using a standard fluorescence-activated cell sorter (FACS, BD Biosciences) together with a similar amount of non-photoactivated control cells. The time between phototagging and FACS sorting was less than 20-30min.

### Single cell proteomics

SCoPE-MS^4^ has been widely adapted and optimized in sample preparation, liquid chromatography and MS settings^18–20^. SCoPE-MS combines the tandem mass tag (TMT) technology with an addition of carrier cells to identify and quantify peptides/proteins of single cells. We prepared the sample using the minimal ProteOmic sample Preparation method (mPOP^21^): a 96-well plate pre-filled with 20 μL pure water and sorted with designated cells per well was frozen on dry ice for 5 min and heated by ThermoMixer C (Eppendorf) at 95 °C for 10 min followed by spinning down at 3000 rpm for 1 min. 20 μL of 100 mM triethylammonium bicarbonate buffer (TEABC, Sigma-Aldrich) was added to each well of the plate, and 1 and 2 μL of 50 ng/μL trypsin (in 100 mM TEABC, Promega) was added to the wells with single cells and two hundred carrier cells, respectively. Digestion was performed at 37 °C ThermoMixer C with shaking speed at 650 rpm overnight. After digestion, the 96-well plate was then spun down at 3000 rpm for 1 min.

0.5 and 1 μL of 85 mM TMT labeling reagent (TMT10plex, Thermo Fischer) was then added to the wells with single cells and two hundred carrier cells, respectively. The labeling was performed at 25 °C with shaking speed of 650 rpm for 1 hr. After labeling, 0.5 μL of 5% (v/v) hydroxylamine was added to each well, and the TMT labeling reaction was quenched at 25 °C with shaking speed at 650 rpm for 15 min. All corresponding samples were combined into the same wells, respectively. 1 μL of 10% (v/v) formic acid (FA, Sigma-Aldrich) was added to each combined well. After acidifying, the samples were desalted by μ-C18 ZipTip (EMD Millipore) and kept in the ZipTip at −80 °C before the MS analysis.

Prior to the MS analysis, the samples were eluted by 50% (v/v) acetonitrile (ACN, Sigma-Aldrich) and speed-vacuum dried. The samples were resuspended with 0.1% (v/v) FA. Nanoflow liquid chromatography tandem mass spectrometry (LC-MS/MS) was performed on an EASY-nLC 1200 (Thermo Fischer) coupled to an Orbitrap Eclipse Tribid mass spectrometer (Thermo Fischer) operating in positive mode. Peptide mixtures were trapped on a 2 cm × 100 μm Pepmap C18 column (Thermo Fisher 164564) and then separated on an in-house packed 50 cm × 75 μm capillary column with 1.9 μm Reprosil-Pur C18 beads (Dr. Maisch) at a flowrate of 250 nL/min, using a linear gradient of 0–32% acetonitrile (in 0.1% formic acid) during 120 min. The MS was performed in the data-dependent acquisition mode. Surveying full scan (MS1) was in the range of 375–1,400 *m/z* and the resolution was set to 120K. Fragmentation of the peptides was performed by HCD. The resolution of tandem mass spectrum (MS2) was set to 30K, automatic gain control (AGC) was 5E4 and the maximum injection time (IT) was 300 ms (Figure S6).

The mass spectrometry proteomics raw data as well as protein and peptide ID lists will be deposited to the ProteomeXchange Consortium (http://proteomecentral.proteomexchange.org).

### Proteomic analysis

Raw MS data were processed with MaxQuant (version 1.6.10.43): peptides were searched against SwissPort database *(Homo sapiens,* downloaded on 2018/12/14), static modification was left empty, variable modifications were deamidation (NQ) and oxidation (M), and minimum peptide length was 5. The reporter ion MS2 analysis was used with the isotopic impurity correction factors provided by the manufacturer (TMT batch number: VB287465). Match between runs was used for identification. Other parameters were remained default. The proteins were filtered at 1% protein identification false discovery rate. Subsequently, the protein groups, peptide list, and peptide-spectrum matches (PSMs) were exported from MaxQuant for further processing. The protein groups list was further imported into Perseus (version 1.6.14.0) for differential protein analysis. The reversed proteins and contaminant proteins were removed, after which 2,001 proteins (14,207 unique peptides) were identified. After that, the cell types (Group 1 or Group 2) were annotated, the intensities were log2-transformed, differential protein analysis by two-tailed t-test was computed, and significantly up-regulated proteins (p ≤ 0.05) were reported and highlighted in the volcano plot (Figure 3c).

**Figure 3.**
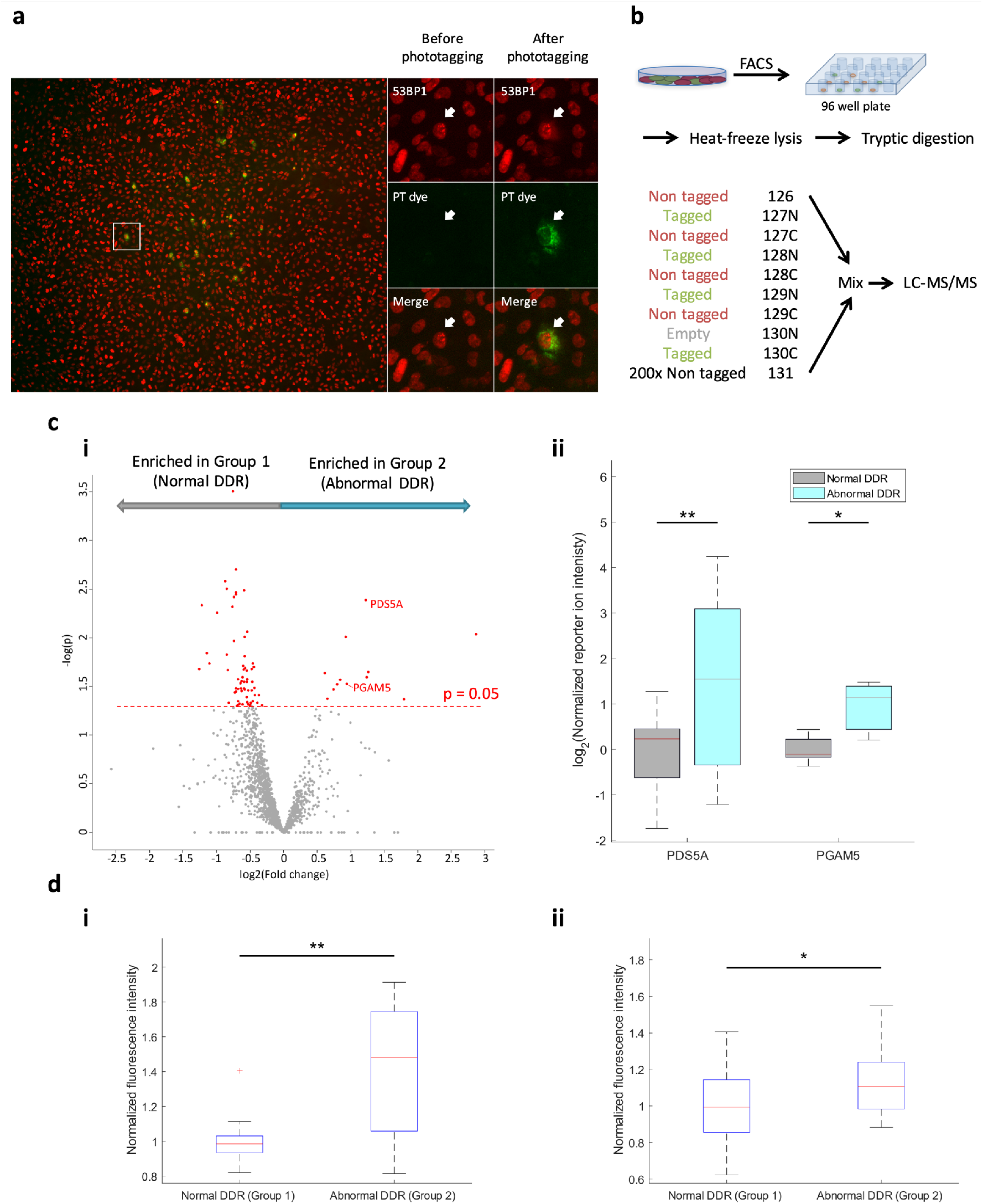
(a) Phototagging of the Group 2 cells. The zoomed-in images showed a representative cell before and after phototagging. Red, 53BP1; green, phototagging (PT) dye. (b) Schematic protocol of single cell proteomics analysis. Four non-tagged cells, four tagged cells and 200 nontagged cells (serving as carrier cells), were labeled with respective 10-plex TMT labels as indicated, and then mixed into one sample before subjecting to LC-MS/MS. 10 samples, in total 40 Group 1 and 40 Group 2 cells, were analyzed. (c) (i) A volcano plot showing proteins enriched either in Group 1 (normal DDR) or Group 2 (abnormal DDR) cells. 80 cells were pooled into the analysis. Dashed red line shows the cutoff of p value at 0.05. (ii) PDS5A and PGAM5 were found upregulated in Group 2 cells and were highlighted. (d) Immunofluorescence quantification of (i) PDS5A and (ii) PGAM5 proteins in Group 1 and Group 2 cells; N = 16 (for PDS5A) and N = 20 (for PGAM5) with p value of 0.0010 and 0.0362, respectively.

### Immunofluorescence staining and analysis

Cells were prepared as the abovementioned cell preparation method. After IR the cells were immediately imaged for 24hr followed by cell fixation with 4% formaldehyde (v/v) in PBS. After fixation the cells were rinsed with PBS containing 0.1% (v/v) Triton followed by incubating with 0.15% (w/w) glycine and 0.5% (w/w) BSA to block non-specific binding sites of the cells. The cells were then incubated with 1:500 rabbit polyclonal anti-PDS5A antibody (Novus), 1:1000 rabbit polyclonal anti-PGAM5 antibody (Novus) or 1:1000 rabbit polyclonal anti-γH2AX antibody (Abcam) for 90 min at room temperature. After antibody incubation the cells were washed with PBS containing 0.1% (v/v) Triton and then incubated with 1:1000 goat polyclonal Alexa-488 anti-rabbit antibody (Abcam) for 60 min at room temperature under dark environment; after that, the cells were then rinsed with PBS containing 0.1% Triton and stored in PBS before imaging. The fixed cells were imaged by a confocal microscope (SP5, Leica); Alexa 488 and mScarlet were excited by the 488 nm and 561 nm lasers, respectively, and imaged by a photomultiplier tube (PMT) with emission spectra setting at 500 nm – 550 nm and 570 nm – 600 nm, respectively. The protein expression level of individual cells was quantified by the summation intensity projection of fluorescence intensity. Group 2 cells were computed against the same amount of randomly selected Group 1 cells and the significance was computed using the two-tailed t-test.

## Supporting information

FUNpro_Supplementary Information

## Acknowledgements

MPC acknowledges support from the Oncode Institute, Cancer GenomiCs.nl (CGC), NWO (the Netherlands Organization for Scientific Research) Veni Grant, Stichting Ammodo and Erasmus MC grant. MPC appreciates Josephine Nefkens Stichting’s support on the UFO microscope. CCH acknowledges support from Ministry of Science and Technology (MOST) in Taiwan (Dragon Gate program: 107-2911-I-002-577 and Columbus Program: 108-2636-M-002-008-&109-2636-M-002-005-). We thank Hanny Odijk for the kind gift of the PB-mScarlet-53BP1 plasmid. We thank Daan Brinks for the discussion and advice on the manuscript.

## Author contributions

PRS conducted the experiments, improved the setup, scripted the algorithm for DDR foci dynamics analysis and analyzed the image analysis data. LY scripted the mTGMM cell tracking algorithm. CB contributed to part of the cell culture preparation for the experiments and the western blot experiment. KB and MP performed the mass spectrometry experiment. RK advised on part of the experimental design and data interpretation. PRS, JD, MP, CCH and MPC advised on and analyzed the single cell proteomic data. MPC and PRS designed most of the experiments. MPC, PRS and CCH wrote the paper with input from all authors. MPC initiated the project. MPC and CCH contributed to and supervised all aspects of the project.

