## Supplementary material for "Functional single cell proteomic profiling of cells with abnormal DNA damage response dynamics": FUNpro_Supplementary Information

### Supplementary Figure S1-S9

**a**

| Algorithm | Precision | Recall | F-score |
|---|---|---|---|
| **Simple Thresholding** | 0.016 | 0.495 | 0.031 |
| **Regional Local Maximum** | 0.482 | 0.448 | 0.464 |
| **Band-pass Filtering** | 0.208 | 0.192 | 0.200 |
| **H-Dome Detection**[1?] | 0.526 | 0.347 | 0.418 |
| **Kernel Density Estimation** | 0.284 | 0.347 | 0.312 |
| **Local Comparison**[2?] | 0.812 | 0.741 | 0.775 |
| **Modified Local Comparison** | 0.935 | 0.727 | 0.818 |
| **Locally Enhancing Filtering**[2?] | 0.466 | 0.370 | 0.413 |
| **Morphometry**[3?] | 0.048 | 0.721 | 0.091 |
| **Multiscale Wavelets**[4?] | 0.026 | 0.865 | 0.051 |

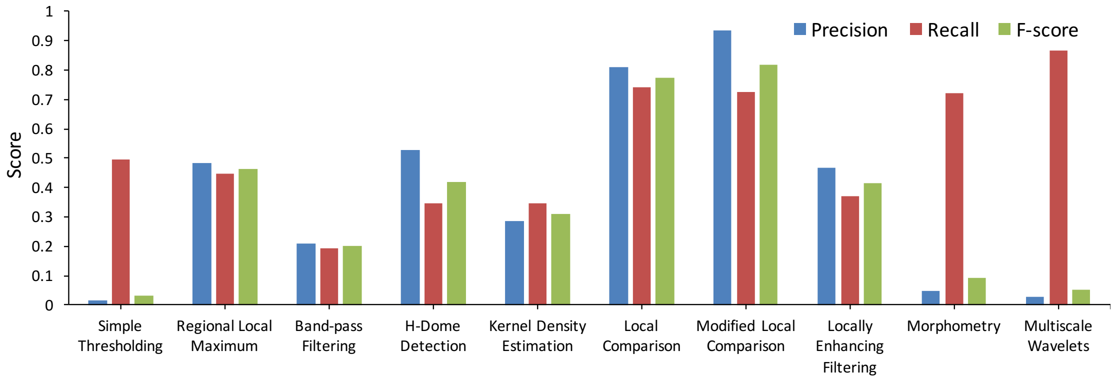


**b**

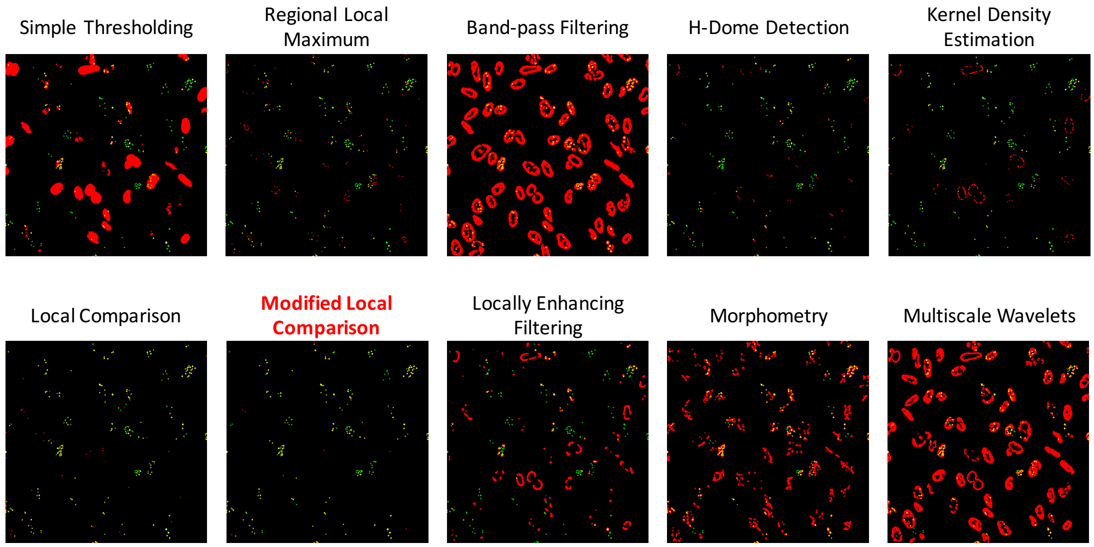


**Figure S1**. (a) Statistics scores of ten different foci detection algorithms. Precision, recall and F score were shown. (b) Images processed by ten different foci detection algorithms with the most optimal conditions. Green: false negative; red: false positive; yellow: true positive.

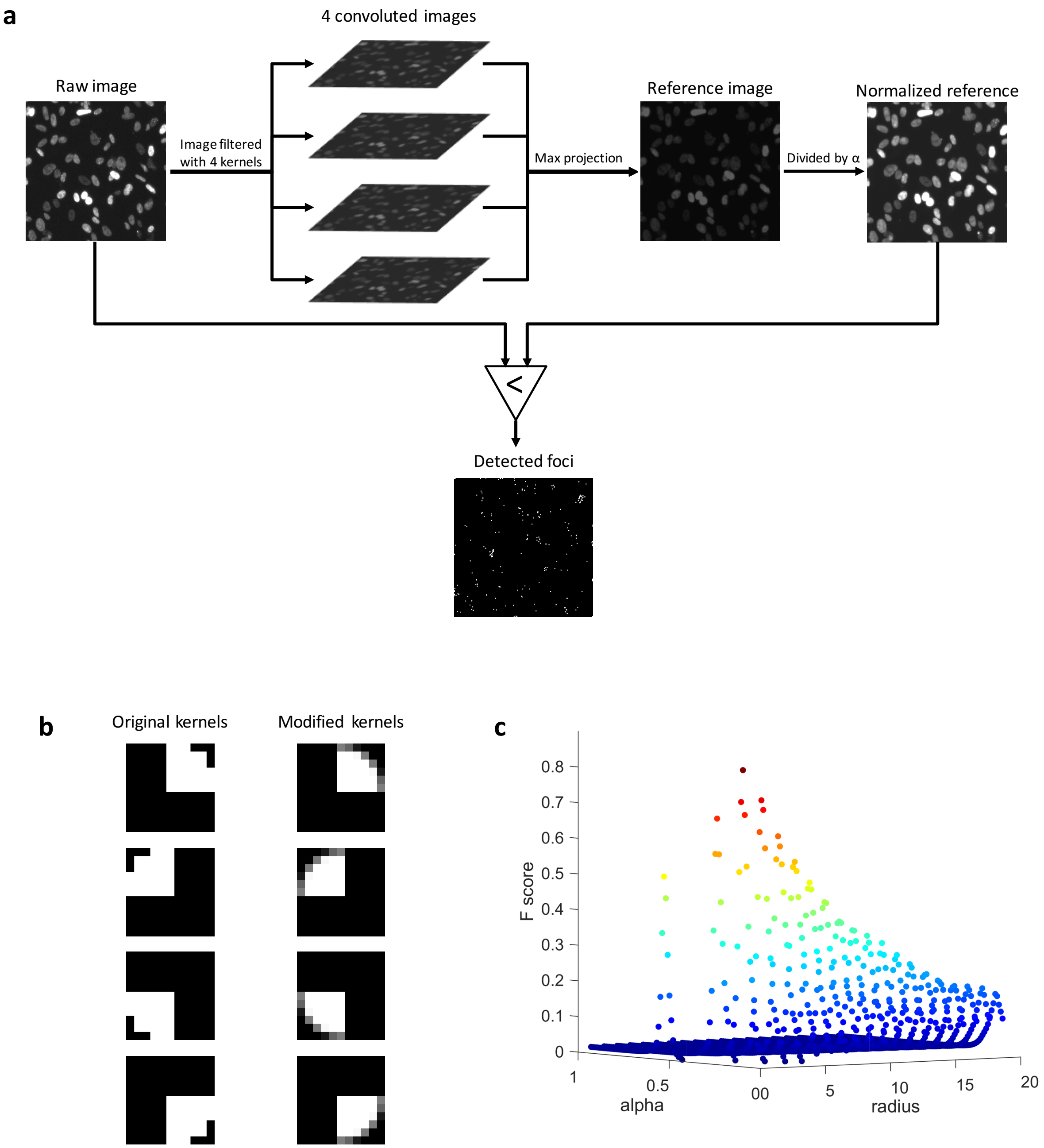


**Figure S2**. Foci detection algorithm. (a) Processing steps of the modified local comparison algorithm. (b) Four original and modified filter kernels used in the modified local comparison algorithm. The modified kernels are Gaussian-like filters. Radius = 5 pixels. (c) The best sensitivity factor α, 0.31, and radius of the filter size, 3 pixels, were optimized based on the highest F score and used in the modified local comparison algorithm for foci detection.

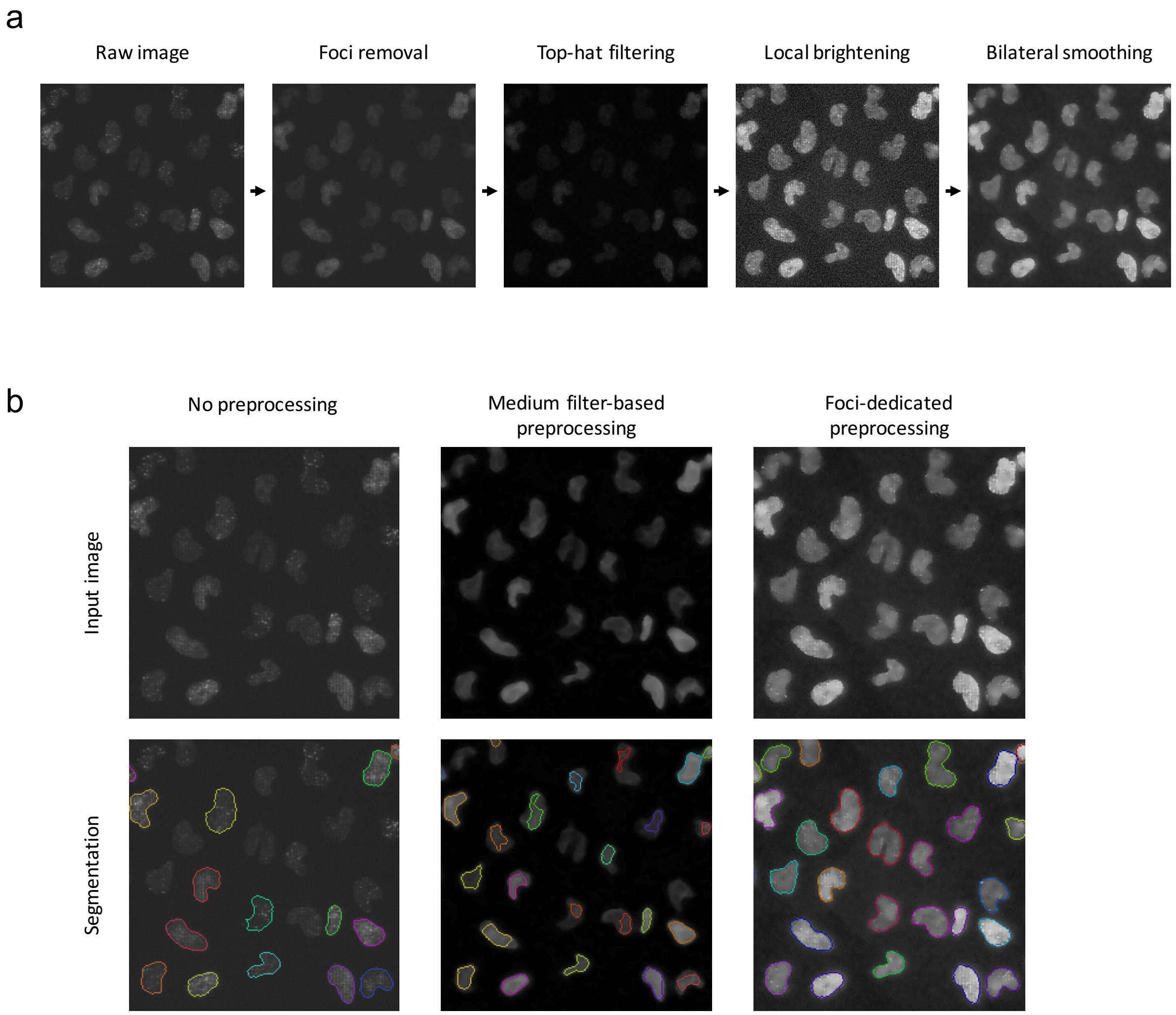


**Figure S3**. Cell (nuclei) segmentation. (a) Processing steps of foci-dedicated preprocessing for nuclei detection. (b) Comparison between medium filter-based vs foci-dedicated preprocessing methods for nuclei detection and segmentation.

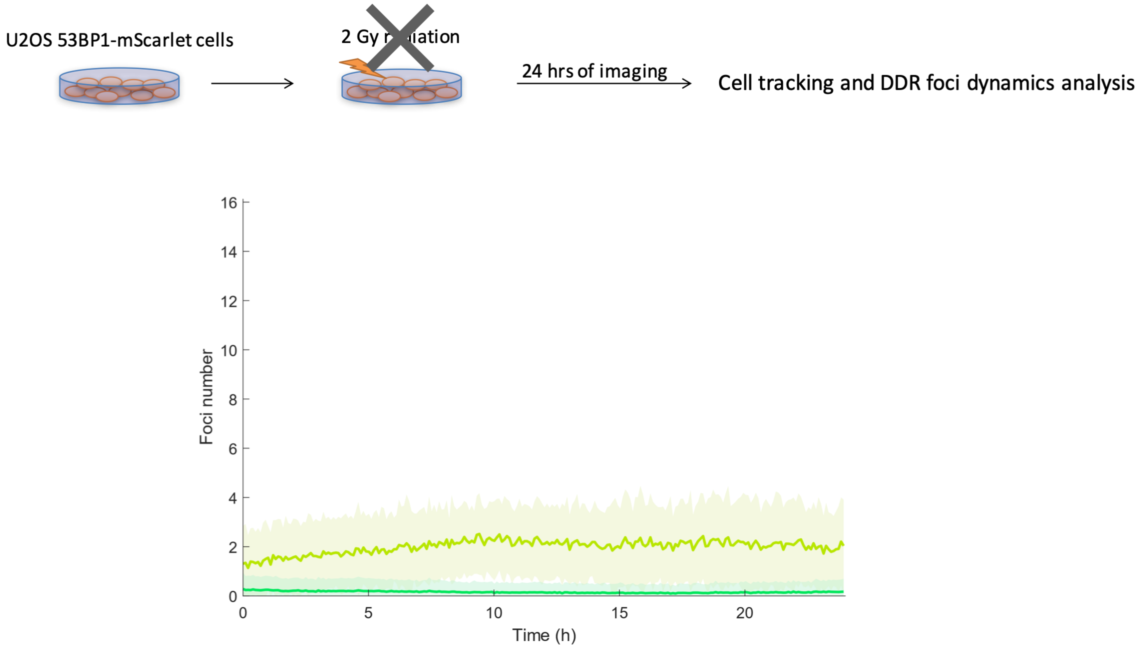


**Figure S4**. 53BP1 foci dynamics of U2OS-53BP1-mScarlet cells over a course of 1 day without irradiation. The foci dynamics trends observed in the Group 1 and Group 2 cells were not seen in this condition. N = 842.

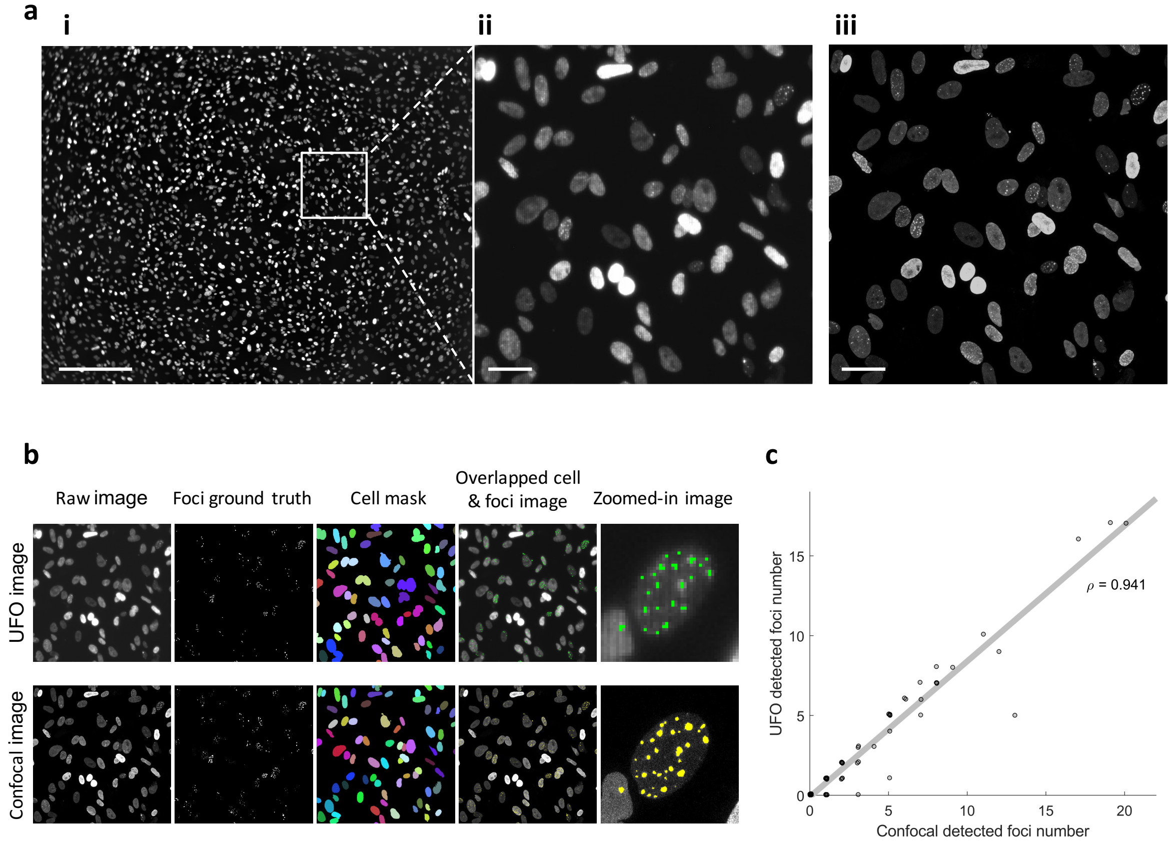


**Figure S5**. DDR Foci detected by the UFO or confocal microscope. (a-i-ii) UFO image of U2OS 53BP1-mScarlet cells (i); scale bar = 500 μm. (ii) A zoomed-in image from (i); scale bar = 50 μm. (a-iii) A representative confocal microscopic image; scale bar = 50 μm. (b) The same field of view of the same dish was imaged under the UFO or confocal microscope. The foci ground truth was manually annotated. (c) Spearman correlation analysis of the foci number per cell between the UFO image and the confocal image (foci diameter larger than 1 μm with SNR larger than 2.0.). Spearman correlation coefficient (ρ) was indicated.

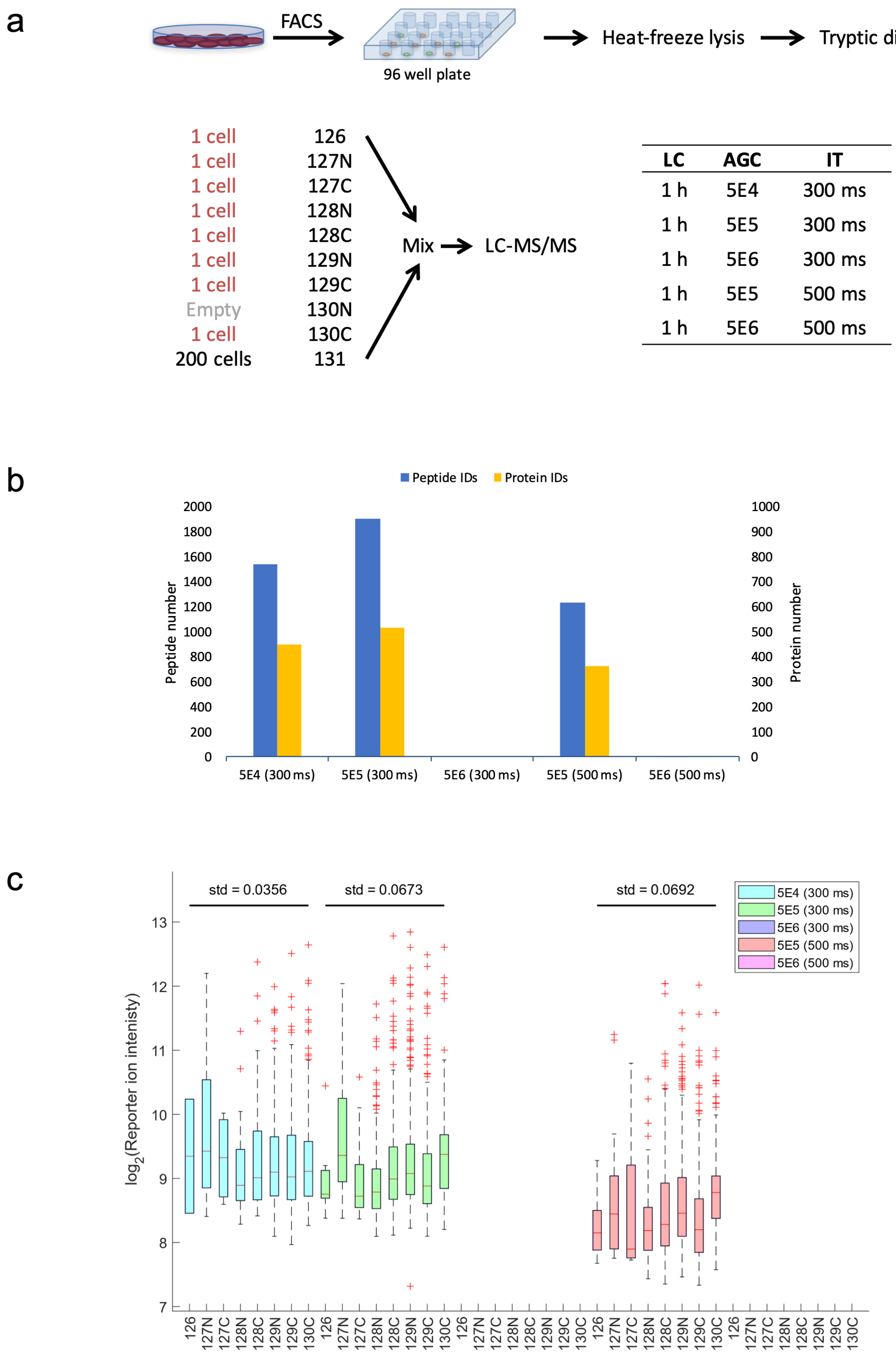


**Figure S6**. Mass spectrometry setting optimization. (a) Schematic of the optimization of automatic gain control (AGC) and injection time (IT). Gradient time for the liquid chromatography is one hour. (b) The total protein and peptide number identified with different AGC and IT settings. (c) Reporter ion intensity distribution of all sample channels. Standard deviations (std) of medians are listed.

a

b

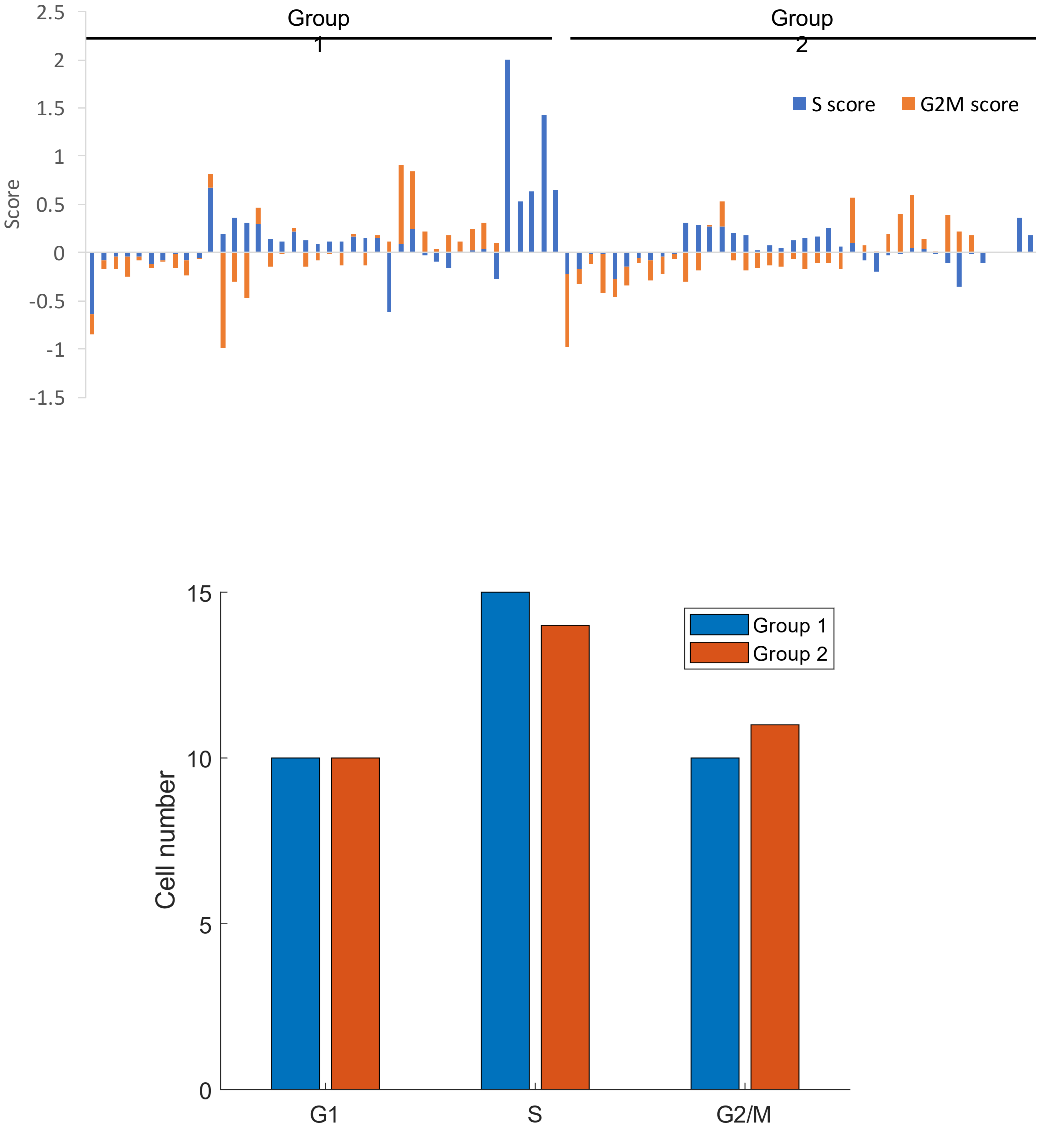


**Figure S7**. Cell cycle scoring analysis. (a) Cell cycle scoring analysis is used to determine cell cycle phase of 80 cells in both Group 1 and Group 2 cells. (b) Cell number of G1, S and G2/M cells from Group 1 (normal DDR) and Group 2 (abnormal DDR) cells.

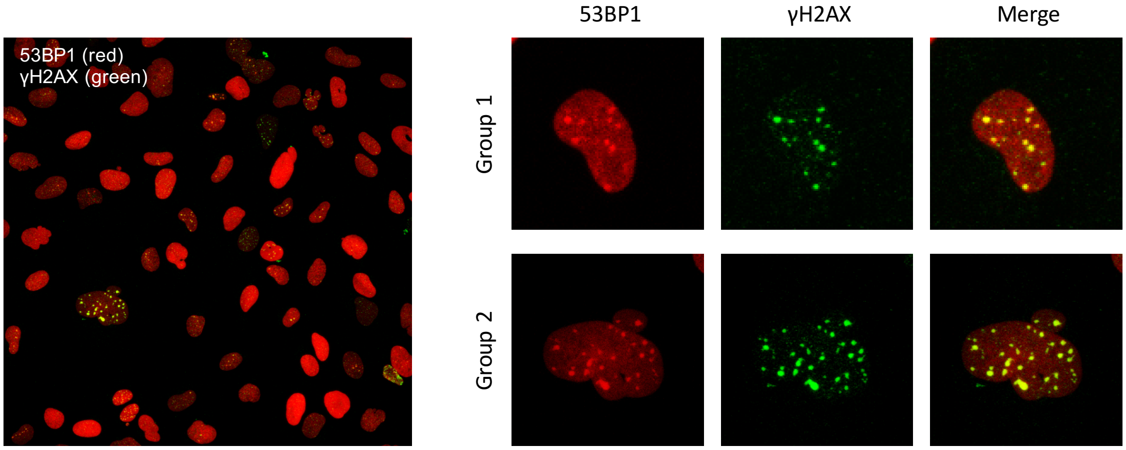


**Figure S8**. A confocal microscopic image (left panel) of U2OS cells expressing 53BP1-mScarlet (red) and immunostained against anti- γH2AX antibodies (green). The cells were imaged after 24 hrs of ionizing radiation. Right panel: One representative zoomed-in image of Group 1 or Group 2 cell.

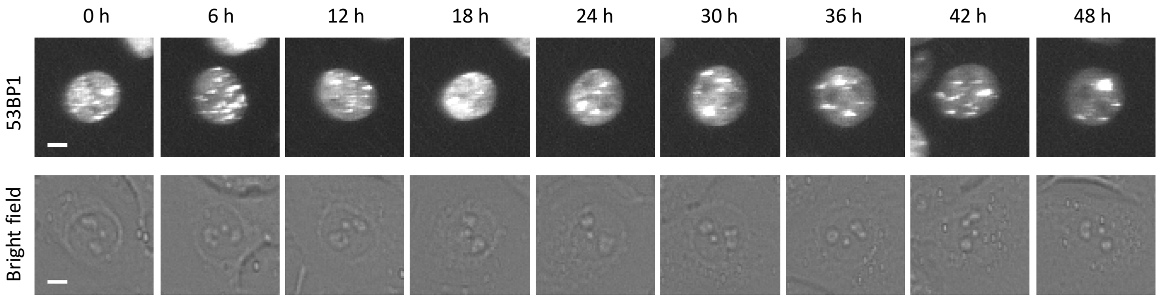


**Figure S9**. Bright field and fluorescent (53BP1) images of one representative Group 2 cell over the course of 2 days after irradiation. Some foci still remain after 2 days of irradiation. Scale bar = 10 μm.
